# A functional overlap between evidence accumulation, confidence, and changes of mind in the pre-supplementary motor area and insula

**DOI:** 10.1101/2024.03.23.586414

**Authors:** Dorian Goueytes, François Stockart, Alexis Robin, Lucien Gyger, Martin Rouy, Dominique Hoffmann, Lorella Minotti, Philippe Kahane, Michael Pereira, Nathan Faivre

## Abstract

Evidence accumulation models have proven to be powerful tools to explain the temporal dynamics of decisions, as well as their metacognitive components such as confidence judgments and changes of mind. However, it is still unclear how and where in the brain evidence accumulation leads to these two metacognitive components. To better understand the functional overlap between evidence accumulation, confidence, and changes of mind, we recorded intracranial high-gamma activity in patients with focal epilepsy while they reported the motion direction of a random dot kinetogram with a computer mouse, and estimated their confidence level. We found an anatomical overlap between the neural correlates of evidence accumulation, confidence, and changes of mind in the pre-supplementary motor area, as well as in the orbitofrontal, inferior frontal, and insular cortices. Both mouse-tracking behaviour and electrophysiological results were reproduced with a post-decisional evidence accumulation model. After characterising the temporal dynamics of decision-making with mouse-tracking and intracranial electrophysiology, we conclude that confidence and changes of mind result from evidence accumulation instantiated before the decision in the pre-supplementary motor area, and after the decision in the insula.

## Introduction

Most of our decisions are accompanied by a feeling of confidence. This feeling, which is thought to derive from metacognitive processes (Yeung & Summerfield, 2012), is essential in a world where immediate feedback rarely follows decisions, and where agents must quickly adjust or even change their decisions based on internal evaluations (Stone et al., 2022). In recent years, evidence accumulation models have been successfully used to describe how decisions unfold over time (O’Connell & Kelly, 2021; Ratcliff et al., 2016; Shadlen & Kiani, 2013), and to understand confidence judgements and changes of mind (CoMs) as part of the decision-making process (Boldt et al., 2019; Pleskac & Busemeyer, 2010; Ratcliff & Starns, 2013; Resulaj et al., 2009). According to *bounded evidence accumulation*, sensory evidence is accumulated until a decision bound is reached (Kiani et al., 2008; O’Connell et al., 2012). Some computational and neurophysiological evidence suggests that the state of the accumulators at the decision time can explain confidence (Kiani et al., 2014; Kiani & Shadlen, 2009). However, there is also compelling evidence that both confidence (Desender et al., 2019; Pereira et al., 2021; Pleskac & Busemeyer, 2010; van den Berg et al., 2016) and changes of mind (Murphy et al., 2015; Resulaj et al., 2009) are well described by models assuming that evidence continues to accumulate after the decision bound is reached (i.e., post-decisional evidence accumulation, see (Desender et al., 2021)). Overall, the nature of the transformation between accumulated evidence and confidence remains debated (Moran et al., 2015; Pleskac & Busemeyer, 2010; van den Berg et al., 2016). Moreover, its neural implementation is also unclear, with numerous discrepancies depending on the model, task, and imaging methods (Balsdon et al., 2021; Dou et al., 2023; Gherman & Philiastides, 2015, 2018; Grogan et al., 2023; Kiani & Shadlen, 2009; Pereira et al., 2021; Pinto et al., 2022). To characterise how decisional and post-decisional processes unfold in time both at the behavioural and neural levels, we combined stereotactic electroencephalography (sEEG) and computational modelling to explore the neural implementation of evidence accumulation, confidence judgments, and changes of mind (CoMs).

We used a protocol to precisely track the decisional and post-decisional temporal dynamics of human participants implanted with sEEG electrodes as they made perceptual decisions. Participants used a computer mouse to respond to a two-alternative forced choice visual discrimination task followed by a confidence rating task. Kinematic analyses of mouse trajectories allowed us to infer both decision times (i.e. when participants started to move the mouse to respond) and CoMs (i.e., when participants modified their trajectory to respond), which we used to temporally realign sEEG data. We harnessed the broad coverage and fine spatial resolution of sEEG to isolate neural markers of evidence accumulation and explored how these markers overlap with markers of confidence and CoMs. Based on an evidence accumulation model, we expected to find early neural correlates of evidence accumulation to determine decision time and accuracy, as well as later correlates to continue after the decision to guide confidence and changes of mind. Together, our results support the co-existence of pre- and post-decisional evidence accumulation in the cortex, and disentangle their relative contribution to decisions, confidence, and changes of mind.

## Results

Twenty-four participants with drug-resistant focal epilepsy performed a two-alternative forced-choice visual discrimination task. They had to indicate whether a random-dot kinetogram was moving toward the right or left side of the screen. Participants reported their choice by moving a cursor with the mouse to click on one of two circles in the top-left and top-right corners of the screen, corresponding to each motion direction (Fig. 1A). Mouse-tracking allowed us to define movement onset as a proxy for decision time and to detect changes of mind and their onsets through changes of trajectories. This visual discrimination task was followed by a confidence judgement where participants were asked to rate their confidence in their initial choice on a continuous scale from 0 to 100 (0: Sure incorrect, 50: Unsure, 100: Sure correct).

**Figure 1.**
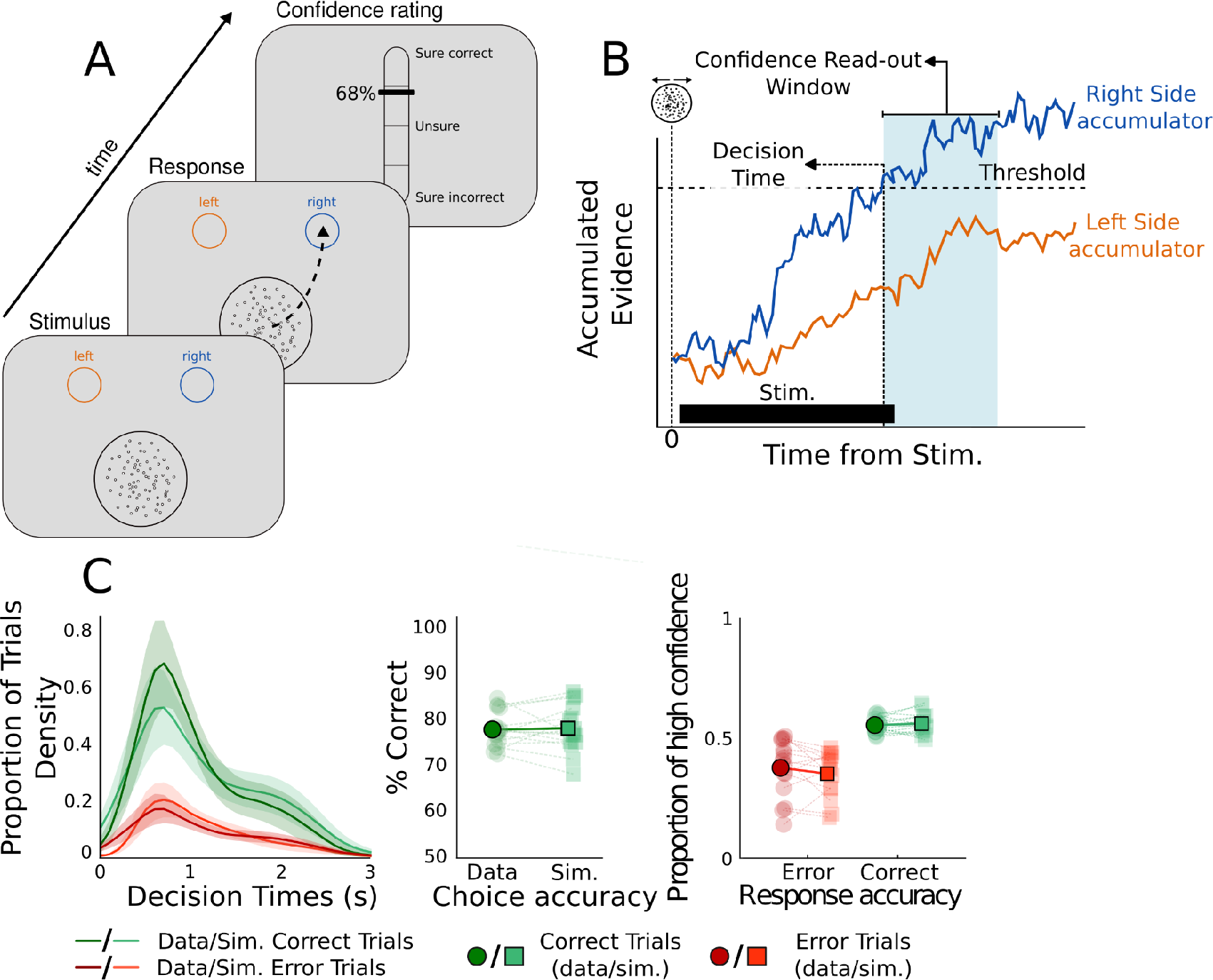
behavioural and modelling results. (**A**) behavioural paradigm. Participants observed the random dot kinetogram and reported its motion direction using the mouse while it remained on screen. Confidence was rated on a continuous scale ranging from 0 to 100. (**B**) Schematic description of the race diffusion model used to fit data. A decision was defined when one of two accumulators coding for each possible motion direction crossed a threshold. Confidence was computed as a read-out of evidence accumulated up to 1s after decision time (confidence read-out window, in blue). (**C**) Comparison of simulated and data performance. The left panel shows the observed and modelled decision time for correct (green) and error (red) trials (shaded areas represent 95% CI). The middle panel shows the observed and modelled response accuracies. The right panel shows the observed and modelled proportions of trials with low and high confidence for correct (green) and error (red) trials. Individual averages are plotted in light green and population averages in dark green.

### behavioural results

All participants were able to perform the task, with an average response accuracy computed at click time of 78.76% (sd = 6.78%), an average confidence judgement of 78.81% (sd = 15.27%), a mean decision time of 1.43s (sd = 1.1s) and a mean response click time of 2.89s (sd = 1.36s). All trials with decision time greater than 2.5s (17.02% of total trials) were excluded from further analyses, as they mainly corresponded to noisy mouse trajectories and unreliable sEEG data. Using generalised linear mixed-effects regressions, we found an effect of response accuracy on decision times (β = -0.2, SE = 0.09, p<0.001) and on confidence (β = 1.84, SE = 0.41, p<0.001), indicating that participants were quicker and more confident in correct vs. incorrect trials. The relationship between confidence and accuracy at decision time was confirmed by computing the area under the receiving operating characteristic curve (AUROC = 0.59, SE = 0.02). Results obtained in individuals with epilepsy were compared with those obtained in healthy controls (see SI for details). We found no significant difference between patients and controls in terms of performance (Mann-Whitney U test, p = 0.60), decision times for correct (Mann-Whitney U test, p = 0.66) and error trials (Mann-Whitney U test, p = 0.91), and proportion of low, mid or high confidence judgments for correct and error trials (Mann-Whitney U test with Benjamini-Hochberg false discovery rate correction, p > 0.05).

### Response accuracy, decision times and confidence as accumulated evidence

To assess whether evidence accumulation is a possible mechanism underlying our behavioural data, we fitted an evidence accumulation model consisting of two anti-correlated accumulators (one for each possible motion direction) to decision times and response accuracy (Faivre et al., 2021; van den Berg et al., 2016). We simulated confidence judgements as a readout of accumulated evidence between 0 and 1s following the initial choice (Fig 1B; see Methods). The model reproduced both decisions (Fig. 1C; S1A-B) and confidence (Fig. 1C; S1C) data well (all R (Spearman) > 0.6, p < 0.017). We further showed that in 13/15 of the participants, models considering post-decisional evidence accumulation had higher log-likelihoods than models considering evidence accumulation until decision time (i.e., accumulation-to-bound). The average fitted confidence readout time was 0.30 s (SE = 0.09) and longer confidence readout times were associated with higher AUROCs in both simulated (R = 0.79, p < 0.001) and observed data (R = 0.73, p = 0.0021), indicating that post-decisional evidence accumulation contributes to accurate confidence judgements. We also fitted the post-decisional evidence accumulation model to our control group data with similar results (Fig. S2). Although we note that other variations in the way confidence is computed have been proposed (Balsdon et al., 2020; Desender et al., 2021; van den Berg et al., 2016), our results indicate that evidence accumulation is a plausible mechanism underlying perceptual decision-making and confidence judgments in this task.

### Identification of task-selective sEEG channels

We recorded from a total of 1138 channels (each channel corresponding to the bipolar referencing of two physical contacts in the brain). We selected 414 channels that were localised in predefined regions of interest (ROIs) corresponding to areas susceptible to instantiate evidence accumulation or to play a role in confidence judgements based on the literature (Balsdon et al., 2021; Fleming et al., 2012, 2018; Liu & Pleskac, 2011; Pereira et al., 2020, 2021). Namely, we selected channels in the visual (caudal medial visual cortex, rostral medial visual cortex, cuneus), parietal (superior parietal cortex, medial superior parietal cortex, dorsal inferior parietal cortex), dorso-lateral prefrontal (dlPFC), inferior frontal (IFC), orbitofrontal (OFC), and insular cortices, as well as the pre-supplementary motor area (pSMA; see SI for corresponding Brodmann areas). We restricted our analyses to stimulus-responsive channels, defined as showing a significant difference between a baseline window (-500ms to -100ms relative to stimulus onset) and a post-stimulus window (200ms up to median decision time) (Mann-Whitney U test, critical p-value 0.05). A total of 102 channels were selected, and all subsequent analyses were performed on this subset (see SI for results from non-selected channels). We focused our analyses on the high-gamma frequency band (70-150Hz), a proxy for local field potentials (Ray et al., 2008). Next, we compared high-gamma activity in correct and incorrect trials. We found 6 channels out of 102 within our ROIs with higher activity following correct vs. incorrect decisions (permutation test, p = 0.030, 1000 permutations), and none with increased high-gamma activity for incorrect trials (see SI). Because of this low number of identified channels related to error processing, we focused the rest of the analysis on correct trials.

### Identification of sEEG channels reflecting evidence accumulation

We aimed to identify channels exhibiting hallmarks of evidence accumulation (Gherman et al., 2023; Kelly & O’Connell, 2015; Kiani et al., 2008; Kim & Shadlen, 1999; Pereira et al., 2021, Stockart et al., 2024). We looked for channels that exhibited a ramping-up of high-gamma activity following stimulus onset, identified by a significant negative correlation between the slope of this ramping-up and decision times across individual trials. To do so, we fitted a linear function to the high-gamma activity for every correct trial from stimulus onset up to the decision time. Then, we selected channels for which slopes were significantly negatively correlated with decision times (Pearson R test, critical p-value 0.05). 41 channels out of 102 exhibited this correlation, exceeding what was expected by chance only (permutation test, p = 0.0009, see methods). Interestingly, channels reflecting evidence accumulation were found across all selected regions of interest (Fig. 2 and SI Table 2). The widespread repartition of evidence accumulation channels corroborates recent results using whole-brain fMRI and intracranial recordings, showing that accumulators are spatially distributed across the brain (Gherman et al., 2023; Morito & Murata, 2022; Stockart et al., 2024).

**Figure 2.**
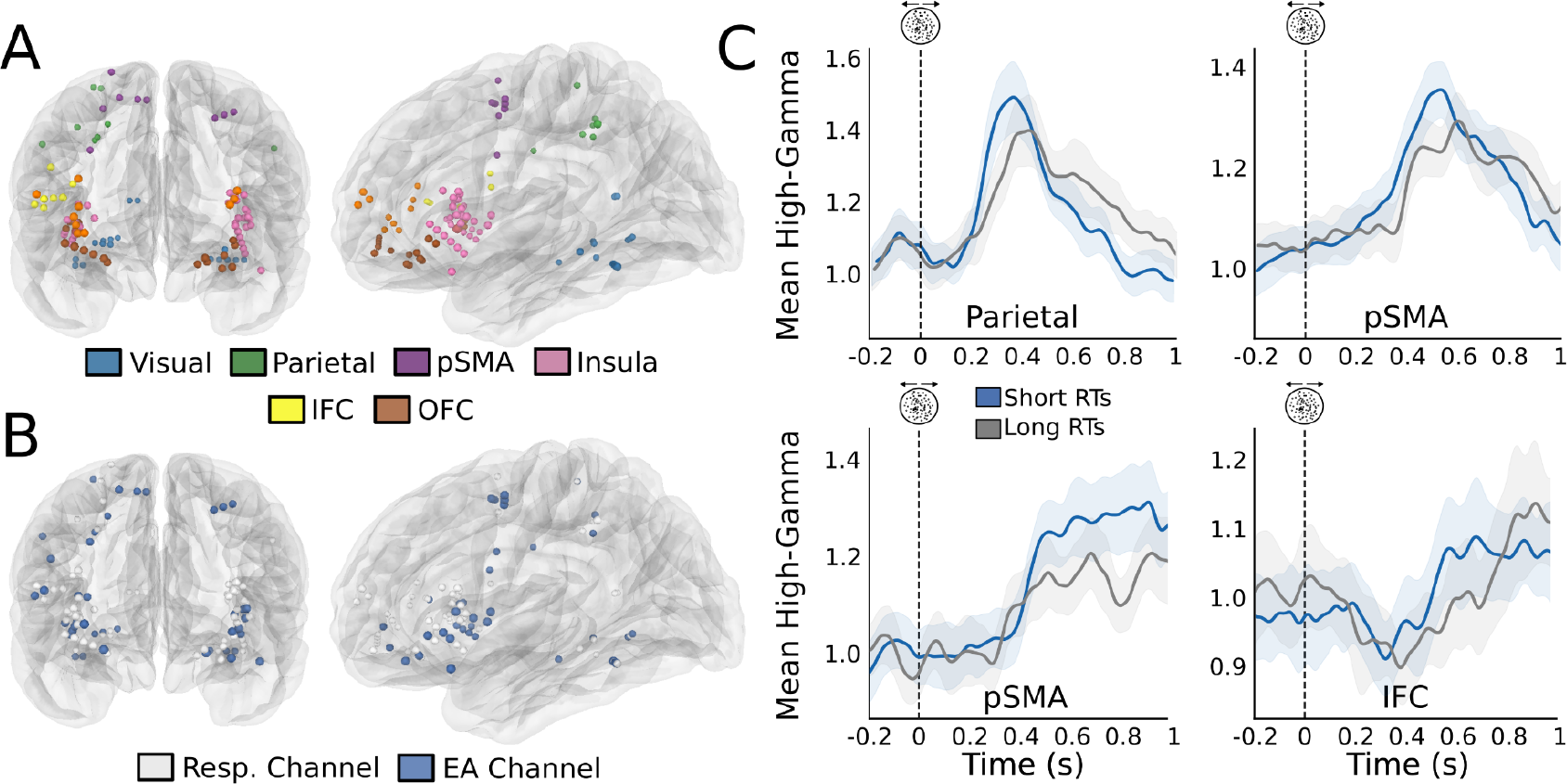
Channels highlighting evidence accumulation in correct trials. (**A**) Channels from all participants are displayed on a template brain based on MNI coordinates. Responsive channels are shown in colours and excluded channels are shown in dark grey. (**B**) Responsive channels reflecting evidence accumulation (blue) or not (white) displayed on the same glass brain as A. (**C**) Examples of channels reflecting evidence accumulation. Each graph represents trial-averaged high-gamma activity aligned on stimulus onset for all correct trials for one selected channel. Even if statistics were performed on continuous decision times, and for illustration purposes only, blue lines correspond to averaged high-gamma for the 50% fastest trials and light grey lines the 50% slowest. Shaded areas represent 95% CI.

### Identification of sEEG channels reflecting confidence

Next, we examined the relationship between single-channel activity before correct decisions and confidence. To do so, we correlated mean high-gamma activity in a 400 ms window preceding decision time with confidence judgments. We found 9 such channels in the visual cortex, parietal cortex and the pSMA, corresponding to 21.95**%** of the evidence accumulation channels (permutation test: p = 0.0019). To support the results of our model indicating that confidence was better predicted when allowing evidence accumulation to continue after the decision, we then aimed to establish if the activity of evidence accumulation channels correlated with confidence following decision time. To do so, we correlated mean high-gamma activity in a 400 ms window following decision time with confidence judgments. We found 11 channels with a positive correlation (Spearman Rank Order correlation test), corresponding to 26.82% of the evidence accumulation channel pool (permutation test: p = 0.0009, SI Table 2) (Fig. 3). Five of these channels (all but one localised in the pSMA and the OFC) also reflected pre-decisional evidence accumulation. Interestingly, the subset of channels reflecting post-decisional confidence was primarily localised in frontal areas (81.81 % of channels, 9 out of 11, 7 in the insula and 2 in the OFC, with the two remaining channels located in the visual cortex). This result shows that a significant proportion of channels exhibiting evidence accumulation-like activity following stimulus onset also co-varied with confidence after the decision, supporting the view that post-decisional evidence accumulation subserves confidence judgements. We also performed a similar analysis including all 102 responsive channels, and found that 10 additional channels reflected confidence preceding decision, and 5 following decision, which were not labelled as accumulating evidence (permutation test on all responsive channels, p = 0.0009, see methods). All of these channels were also located in the OFC (2 channels) and the insula (3 channels; see Fig. 3 and SI). As decision time and confidence are correlated (Baranski & Petrusic, 1998; Faivre et al., 2020), we verified that the magnitude of the correlation between high-gamma activity and decision time was significantly lower than that of the correlation between high-gamma activity and confidence (see SI).

**Figure 3.**
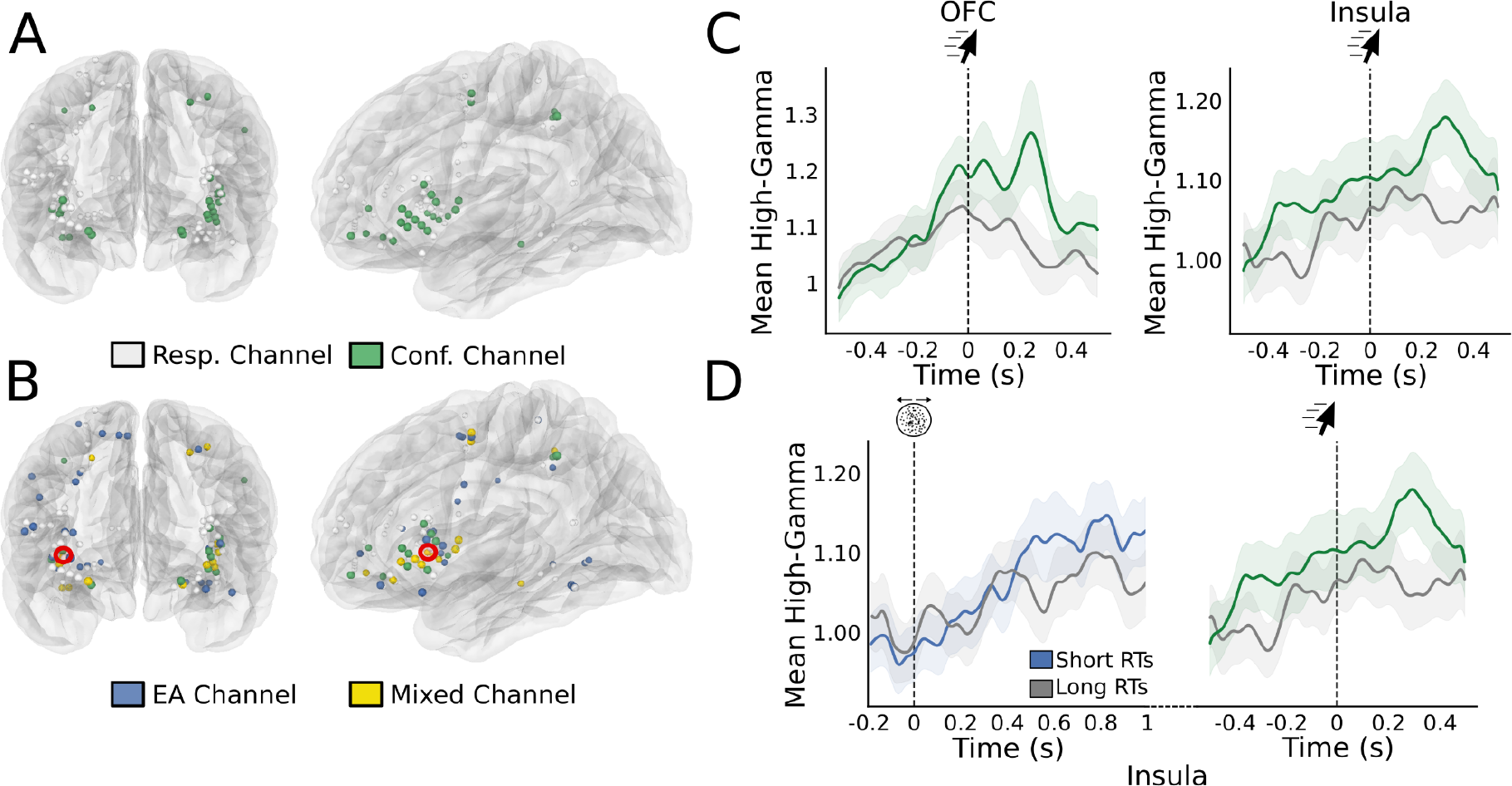
Repartition of channels reflecting evidence accumulation and confidence in correct trials. (**A**) Confidence-sensitive channels plotted in green against all responsive channels in white (**B**) representation of responsive channel (white), evidence accumulation channels (blue), confidence channels (green) and channels with an overlap between evidence accumulation and confidence (yellow). (**C**) Example of two confidence-sensitive channels. Trial-averaged high-gamma activity is plotted aligned on decision time. Green lines correspond to the 50% of trials with the highest confidence and grey to the 50% with the lowest confidence. Shaded areas correspond to 95% CI. (**D**) Example of a channel (highlighted in red in panel B) with an overlap between evidence accumulation and confidence. The left panel corresponds to stimulus-aligned, trial-averaged high-gamma separated based on decision time. The right panel corresponds to movement-aligned high-gamma activity separated based on confidence. Shaded areas correspond to 95% CI.

### Analysis of regions of interest

To substantiate the single-channel analyses and characterise neural activity at the level of cortical regions, we pooled all evidence accumulation channels across all participants for each ROI. Assuming that high-gamma activity related to evidence accumulation ramps up steeper for earlier decision times, we expected to find a relationship between high gamma and decision time after stimulus onset and before decision time (Steinemann et al., 2022). Interestingly, this analysis at the ROI level brushes a more nuanced picture of the anatomical distribution of evidence accumulation, as visual (8 EA-channels), parietal (4 EA-channels), and orbitofrontal (4 EA-channels) areas did not exhibit a significant effect at the ROI level (see Table 1). Only the pSMA (7 EA-channels), the IFC (6 EA-channels), and insula (11 EA-channels) qualified as evidence accumulation ROIs (Fig. 4). Examining the time course of regression coefficients between high-gamma activity and decision times revealed that the insula and the pSMA had similar peak effect latencies after the stimulus onset (pSMA: 537ms, insula: 553ms; Fig. 4.D). However, significant regression coefficients were maximal before the decision in the pSMA, while they were observed after the decision in the insula. These temporal profiles are reminiscent of early and late accumulation profiles (Morito & Murata, 2022; Msheik et al., 2022).

**Figure. 4.**
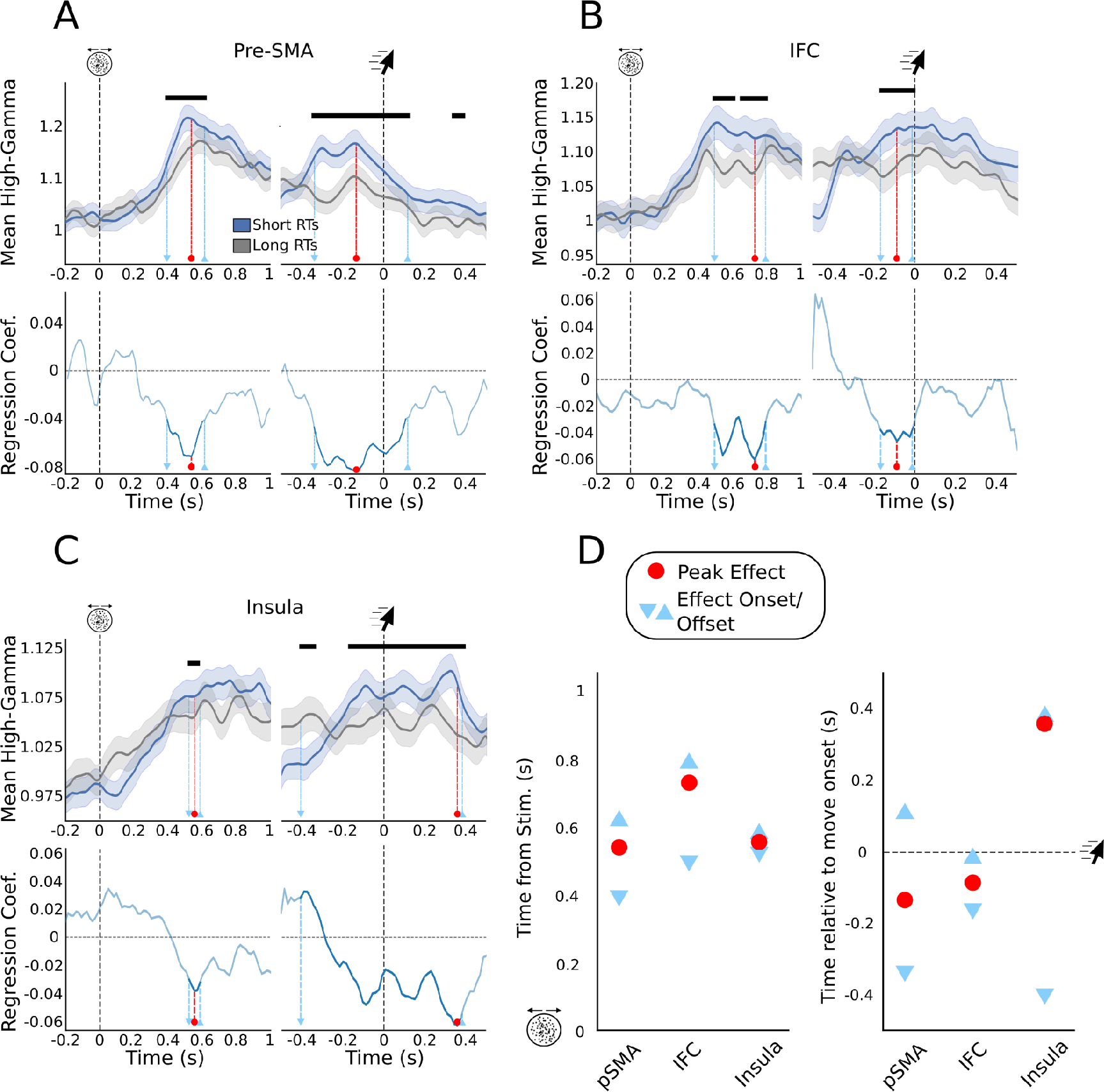
Stimulus-aligned and movement-aligned high-gamma activity for regions of interest reflecting evidence accumulation (**A, B, C**) Top: Stimulus-aligned and movement-aligned high-gamma activity averaged across participants for all EA channels. Dark bars indicate a significant relationship between high-gamma and move onset timing (p<0.05, FDR-corrected). Blue lines correspond to averaged high-gamma for the 50% fastest trials and grey lines for the 50% slowest. Shaded areas correspond to 95% CI. Bottom: Regression coefficient as a function of time in the stimulus- and movement-aligned window. Light blue markers indicate the onset and offset times of significant effect, and the red marker indicates peak effects. (**D**) Latency analysis in the stimulus and move-aligned window for the three ROIs reflecting evidence accumulation. Conventions are similar to panels A to C.

**Table 1:** Summary of ROI analyses.

|  | Evidence-accumulation |  |  | Confidence |  |  | CoM |
| --- | --- | --- | --- | --- | --- | --- | --- |
|  | Single-channel analysis | ROI analysis |  | Single-channel analysis | ROI analysis |  | ROI analysis |
|  |  | EA | Post-decisional effect |  | Confidence | Post-decisional effect |  |
| Visual Cortex | 8 Channels | No | No | 2 Channels | No | No | No |
| Parietal Cortex | 4 Channels | No | No | 3 Channel | **Yes** | No | No |
| pSMA | 7 Channels | **Yes** | **Yes** | 2 Channel | **Yes** | No | **Yes** |
| dIPFC | 1 Channel | No | No | 1 Channel | No | No | **Yes** |
| IFC | 6 Channels | **Yes** | **Yes** | 0 Channel | No | No | No |
| OFC | 4 Channels | No | No | 6 Channels | **Yes** | **Yes** | **Yes** |
| Insula | 11 Channels | **Yes** | **Yes** | 14 Channels | **Yes** | **Yes** | **Yes** |

We used a similar approach to explore the relationship between high-gamma activity and confidence at the ROI level. We assessed if the pooled high-gamma activity of channels reflecting evidence accumulation in each ROI also covaried with confidence after the decision time, similar to what we observed with single channels. Among ROIs reflecting confidence (see Table 1), two different temporal profiles emerged. The first group of areas, including the parietal cortex and the pSMA exhibited early peak effect latencies (average peak latency -246ms) with short-lived effects disappearing before decision time (Fig. 5A, SI. 3A). By contrast, the orbitofrontal and insular cortices showed later peak effect latencies (average -134ms after decision time), with long-lasting effects extending well beyond decision time (Fig. 5B, C, SI. 3B). Although only the pSMA and insula exhibited both markers of evidence accumulation and confidence at the ROI level, a majority of regions with channels accumulating evidence also correlated with confidence. To ensure that these effects were not due to the interaction between confidence and decision time, we ran a similar mixed-effects regression including an interaction term between confidence and decision time (see SI). This model yielded similar results as the ones described above, with minimal interaction effects. This indicates that the effects of confidence on high-gamma activity reported above are not driven by decision time. Finally, to assess if our computational model could account for these temporal dynamics, we simulated traces of evidence accumulation using the individual parameters found to best reproduce our behavioural data (see above). By manipulating the window during which evidence accumulation could take place following decision time, we checked if the different profiles observed in our neural data could be the results of distinct pre- and post-decisional evidence accumulation processes (see methods). Limiting the duration of post-decisional evidence accumulation to the confidence readout time, we could reproduce early/transient correlates of confidence similar to the ones we found in the pSMA. By contrast, allowing post-decisional accumulation for longer durations (500 ms post-decision) resulted in traces resembling the late and sustained correlates of confidence we found in the insula (Fig. 5A/B bottom panels).

**Figure. 5.**
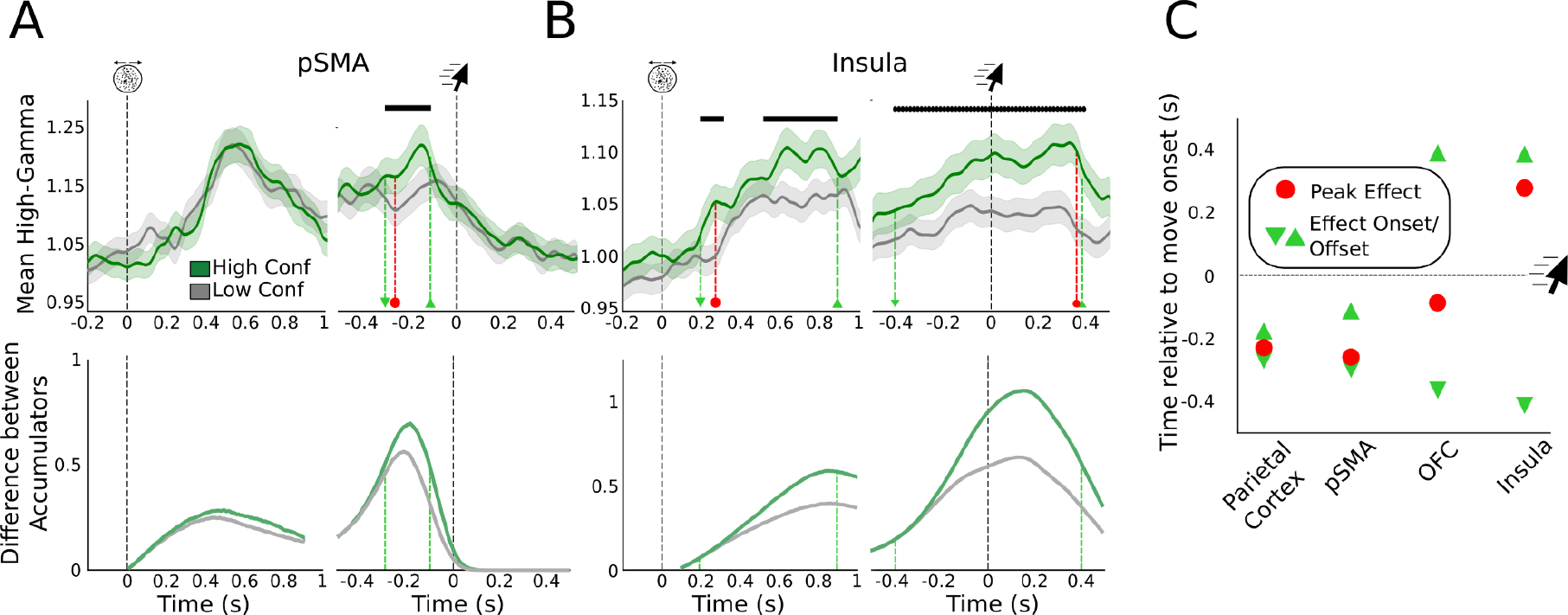
Stimulus-aligned and movement-aligned high-gamma activity in the pSMA and insula as a function of confidence. (**A and B**) Top: Stimulus-aligned and movement-aligned high-gamma activity averaged across participants, including only channels reflecting evidence accumulation in each ROI. Dark bars indicate a significant relationship between high-gamma and confidence (p<0.05, FDR corrected using the Benjamini-Hochberg method). Green lines correspond to averaged high-gamma for the 50% of trials with the highest confidence and grey lines for the 50% lowest. Shaded areas correspond to 95% CI. Dashed light green markers indicate the onset and offset times of significant effects, and dashed red markers indicate peak effects. Bottom: Average difference between accumulator traces simulated by the evidence accumulation model. (**C**) Latency analysis in the movement-aligned window for the four regions sensitive to confidence. Conventions are similar to panels D in Fig. 4.

### Behavioural and neural markers of changes of mind

Having established a link between confidence and post-decisional evidence accumulation, we used a similar approach to explain another manifestation of metacognitive monitoring: changes of mind (CoMs). Indeed, CoMs have been proposed to depend on confidence (van den Berg et al., 2016) or post-decisional evidence accumulation (Burk et al., 2014; Resulaj et al., 2009; van den Berg et al., 2016). Harnessing mouse-tracking, we defined changes of mind as trials in which sudden changes in mouse trajectories occurred before the final response was provided (9.64% of trials, sd = 5.88%, Fig. 6A). Using the same post-decisional evidence accumulation model as for confidence, we simulated the onset of CoMs as the time following the decision when the difference between the two accumulators reached a threshold. By fitting this threshold to the behavioural data of individual participants, our model could predict the distribution of CoM onset timing (Fig. 6B, SI. 1D). Turning to sEEG data, we compared high-gamma activity aligned on CoM onset to the activity of evidence accumulation channels aligned on the initial decision time. We reasoned that if changes of mind involve evidence accumulation, high-gamma activity aligned on CoM onset should closely resemble the activity observed around the initial decision time in channels reflecting evidence accumulation. Thus, we defined an ROI as reflecting CoM if its activity was (1) ramping up in the interval leading to CoM and different from similarly aligned activity in non-CoM trials (CoM-like time) and (2) showed a similar pattern of high-gamma activity leading to CoM in CoM trials and to move onset in non-CoM trials. Such ROIs included the parietal cortex, pSMA, OFC, and insula (Fig.6, Table 1, Table S2). Interestingly, these overlapped both with ROIs reflecting evidence accumulation and ROIs reflecting confidence. We note, however, that the same analysis restricted to channels reflecting evidence accumulation did not yield any significant result, possibly due to lack of statistical power as CoMs were rare.

**Figure 6.**
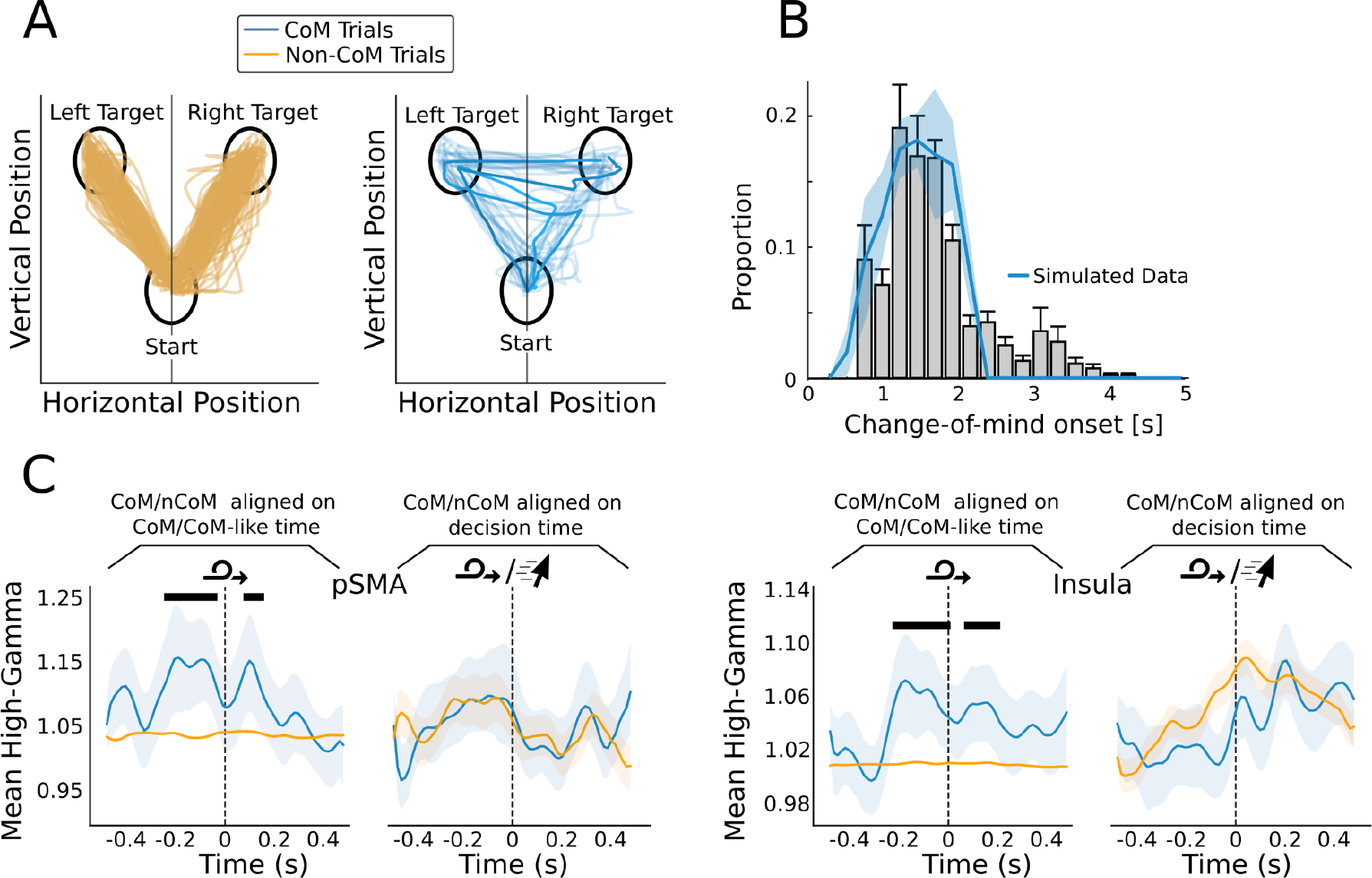
Neural Correlates of Changes of Mind. (**A**) Mouse trajectories for regular trials (left) and CoM trials (right) for one example participant. (**B**) Comparison of observed and simulated CoM onset. Blue line shows simulated data. Grey histogram corresponds to observed CoM-onsets binned and averaged across subjects. (**C**) CoM-aligned high-gamma activity for CoM trials compared to high-gamma aligned on CoM-like times for non-CoM trials (left panel) or move-aligned high-gamma for non-CoM trials (right panel). Dark bars indicate a significant effect of condition (CoM/non-CoM) on high-gamma levels (p<0.05, FDR corrected using the Benjamini-Hochberg method). Shaded areas correspond to 95% CI.

## Discussion

We studied how evidence accumulation underlies confidence and changes of mind during perceptual decision-making. To this end, we fitted a computational model of competing accumulators to human participants’ decisions, response times, and confidence ratings. Examining how intracranial recordings reflected these behavioural variables, we found overlapping regional markers of post-decisional evidence accumulation and confidence in the pSMA and the insula.

### Evidence accumulation and confidence

Activity in the pSMA reflected confidence following stimulus onset until the decision time, in line with a possible role of the pSMA in tracking the uncertainty of a task as it unfolds until a decision is reached (Aquino et al., 2023; Bonini et al., 2014; Fleming et al., 2018; Nachev et al., 2008; Ullsperger et al., 2014). This mechanism may be useful in situations where one needs to adapt behaviour before a decision, for instance when deciding to opt-out before the decision is taken if it is deemed too difficult (Kepecs et al., 2008; Kiani & Shadlen, 2009). The dynamic we observed in the pSMA is consistent with evidence accumulation stopping in the vicinity of decision time (O’Connell et al., 2012; Shadlen & Kiani, 2013). Within such a framework, confidence is derived from the state of the losing accumulator, as by definition the state of the winning accumulator is equal to the bound (Kiani & Shadlen, 2009). The higher the state of the losing accumulator (i.e., the closer it is to the bound), the lower the confidence. Importantly, in all but two participants, an accumulation-to-bound model could not reproduce our confidence data as well as a model reading out confidence after post-decisional evidence accumulation. Furthermore, increased post-decisional time also explained the better metacognitive sensitivity of confidence ratings across participants. Therefore, more than accumulation-to-bound was needed to explain confidence in this task, including post-decisional evidence accumulation up to an early confidence readout latency shortly after decision time (Desender et al., 2021; Pereira et al., 2020; Pleskac & Busemeyer, 2010) consistent with pSMA activity (Figure 5A, Figure S6).

As the state of each of the two accumulators could not be distinguished at the sEEG level (i.e., we could not find univariate or multivariate signatures of rightward vs leftward visual motion), it is unclear what feature of evidence accumulation was reflected by high gamma activity observed in the pSMA. In case it reflected the sum of the two accumulators, high activity would correspond to a high state of the losing accumulator and therefore confidence and high gamma activity should be negatively correlated. This is inconsistent with our results, as we observed a positive correlation between high-gamma activity and confidence.

Instead, a possibility is that high-gamma activity in the pSMA represents the difference between the two accumulators, as was found in the parietal and frontal cortex of rats (Scott et al., 2017). This difference should be large for correct decisions with high confidence, and small for correct decisions with low confidence, which is what we observed in both our data and computational model. According to this hypothesis, confidence depends mostly on the negative evidence encoded by the losing accumulator (Kiani & Shadlen, 2009), which goes against another line of research showing that confidence neglects negative evidence (Peters et al., 2017; Samaha & Denison, 2022; Zylberberg et al., 2012). Interestingly, we could not confirm that activity in the pSMA increased after errors (Fig. S5), despite numerous studies suggesting a role of the pSMA in error monitoring (Bonini et al., 2014; Fu et al., 2019; Scangos et al., 2013). One possible reason is that our task included no response conflict and produced long response times, contrary to what is commonly used in experiments showing increased activity in the pSMA following errors (Bonini et al., 2014; Cavanagh et al., 2012; Fu et al., 2019). Our computational model adds support to this account as it simulated lower activity for errors. According to this account, incorrect trials are similar to low-confidence correct trials with small differences between the losing and winning accumulators.

We also found positive correlations between confidence and high-gamma activity in the insula. In contrast to our findings in the pSMA, correlates of confidence in the insula also extended until after the decision. Our computational model fitted on behavioural data could reproduce qualitatively similar traces when evidence accumulation was allowed to continue after the decision and after the estimated confidence readout time. This indicates that high-gamma activity measured in the insula reflected the difference between the two accumulators following post-decisional processing. This anatomical dissociation between pre- and post-decisional evidence accumulation was further supported by univariate analyses: channels instantiating evidence accumulation also encoded pre-decisional confidence in the pSMA and post-decisional confidence in the insula. Interestingly, in the context of decision-making, stimulating the insula has been found to have an effect on confidence (Cecchi et al., 2023). The insula has also been hypothesised to provide inputs to dorsomedial regions including the pSMA (Bastin et al., 2017), as well as to the OFC (Dhar et al., 2011; Pouget et al., 2016), another region found to reflect confidence in our own data. In addition, contrary to what we found in the pSMA, high-gamma activity in the insula and the OFC was higher for correct than incorrect trials, compatible with the view that these regions are also involved in error monitoring (Fig. S6).

### Evidence accumulation and changes of mind

Post-decisional evidence accumulation was particularly relevant in our task, as perceptual information was still present on screen following the initial decision and participants had some time to change their mind. In other words, even after participants initiated a mouse movement towards one of the two targets, they could consider additional sensory evidence and revise their initial decision to reorient the mouse towards the other target, which they did in about 10% of trials (Resulaj et al., 2009; van den Berg et al., 2016). Overall, the effect of changes of mind on accuracy and confidence were similar to what had already been reported using the same task in a cohort of healthy participants and participants with schizophrenia (Faivre et al., 2021) see SI). Interestingly, we could predict the timing at which such changes of mind occurred using our computational model when it allowed evidence accumulation to continue after the decision time. In line with this modelling result, we found that neural activity in the insula - which reflected post-decisional evidence accumulation and confidence - also reflected the occurrence of a change of mind. Namely, the neural activity time-locked to the onset of changes of mind resembled the activity time-locked to decision-time in trials without changes of mind, suggesting that participants changed their minds by initiating a new decision following post-decisional evidence accumulation. This hypothesis is in line with earlier works indicating that continued processing of information following the initial decision, either due to processing delays or the arrival of new information, may be responsible for the revision of the initial decision (Kiani et al., 2014; Resulaj et al., 2009; van den Berg et al., 2016). We note that these results must be considered cautiously due to the relatively low number of CoM trials. This limitation prevented us from reliably identifying the hallmarks of post-decisional evidence accumulation in trials with changes of mind as we did for initial evidence accumulation in trials with no changes of mind. As a result, factors other than evidence accumulation might have contributed to the difference in high gamma activity between trials with and without changes of mind. Such factors include sensorimotor signalling, as the onset of changes of mind coincided with deviations in mouse trajectories. Importantly, we accounted for motor contributions to our results by re-running population analyses for evidence accumulation and confidence while including instantaneous mouse velocity as a covariate of no-interest, with no impact on our main results (see SI). We also note that motor aspects should not be treated only as a nuisance, as some are considered key factors of decision-making and metacognitive monitoring (Burk et al., 2014; Faivre et al., 2020; Filevich et al., 2020; Fleming et al., 2015; Pereira et al., 2020; Sanchez et al., 2023). Due to this, we are confident that evidence accumulation drives changes of mind in the current task as was shown in several similar tasks (Kiani et al., 2014; Resulaj et al., 2009; van den Berg et al., 2016). Additionally, an alternative account of changes of mind proposes that observers may revise their decisions through feedback signalling taking place at a decisional level (Atiya et al., 2020). Interestingly, we found that activity in the pSMA, which mostly reflected pre-decisional evidence accumulation also distinguished trials with and without changes of mind, further supporting this hypothesis.

### Single channel level vs regional functional markers

A particularity of our analyses was to combine single-channel analyses with a larger-scale approach, which involved mixed-effects regression models combining all channels within a region of interest. Although these two approaches are mutually informative, we found that the distributions of channels reflecting evidence accumulation and confidence do not perfectly overlap with the results obtained at the regional scale. These discrepancies may be simply due to the intrinsic limitations of working in a clinical setting with no control over recording sites. However, it could also indicate that the granularity of accumulators’ distribution may be finer than the spatial extent of the regions of interest we defined. Indeed results from single-unit recordings in non-human primates show that neuronal activity is very heterogeneous even within a small area such as in the lateral intraparietal area (Kiani & Shadlen, 2009; So & Shadlen, 2022). Similarly, spatial selectivity in sEEG was found for language or attention (Flinker et al., 2011; Slama et al., 2021). This possibility would explain the diversity of results observed in the literature when comparing imaging techniques with lower spatial resolution such as EEG, and works using high spatial resolution techniques such as single-unit recordings (Kiani et al., 2008; Morito & Murata, 2022; van Vugt et al., 2012). Finally, it would explain why we did not find correlates of evidence accumulation at the population level in the parietal cortex, despite considerable literature showing its involvement in evidence accumulation processes (Kiani et al., 2008; Platt & Glimcher, 1999; Shadlen & Newsome, 2001). Finally, we could not find a common population code between evidence accumulation and changes of mind using multivariate analyses.

Our results show how evidence accumulation is implemented in the cortex on a global scale, in line with recent results indicating that it is an “ubiquitous” process (Gherman et al., 2023; Morito & Murata, 2022; Msheik et al., 2022). Perhaps more importantly, by combining our modelling and electrophysiological results, we uncovered what could be a functional hierarchy of areas supporting confidence. Indeed, we found regions in which evidence accumulation subserves confidence, either in a pre-decisional manner (e.g., pSMA) or a post-decisional (e.g., insula) manner, as well as regions that did not reflect evidence accumulation but still reflected confidence and changes of mind (e.g., OFC). These regions might thus integrate the activity of regions exhibiting evidence accumulation and serve as output systems. Overall, our results provide neurophysiological evidence to explain how a hierarchy of accumulators could provide a diversity of metacognitive processes unfolding at different moments in time.

## Material and Methods

### Participants

24 individuals with pharmacologically intractable epilepsy undergoing treatment at the University Hospital in Grenoble, France were recorded, among which 21 were included in the present article (8 females, mean age = 31.90, sd = 9.78, see Table 3 SI for details). Patients were implanted with semi-rigid linear electrodes in preparation for the surgical resection of the seizure focus. The experimental protocol was approved by an ethical committee (MapCog_SEEG 2017-A03248-45), and informed consent was obtained from each patient. The electrode locations were decided following pre-surgical MRI and were based solely on clinical criteria. The recording sites varied across patients, with an average number of 138.8 channels (sd = 27.87). The experiment was performed on a laptop while the participants sat in bed. One participant was excluded due to missing experimental triggers, and two due to excessive epileptic activity. Four additional participants were excluded from the confidence-related analyses due to a failure to correctly use the confidence scale. They were however included in analyses focusing only on evidence accumulation. 14 participants with more than 10 CoM trials were included in the analyses related to changes of mind.

### Procedure

The task consisted of a random-dot kinematogram 2-AFC discrimination task followed by a confidence judgement. It was implemented under Matlab R2019b using Psychtoolbox-3 (Brainard, 1997; Pelli, 1997) and ran on a Dell Precision 7550 laptop under Ubuntu 20.04. The task was performed at the hospital with the participants sitting in their beds and the laptop placed on a tablet in front of them. In the visual discrimination task, participants were asked to decide whether a cloud of dots circumscribed to a 108 px radius circle was moving rightward or leftward. Responses were given by moving a computer mouse from its initial position at the bottom centre of the screen to one of two targets presented in the top right and left corners of the screen (108 px radius), corresponding to right and left decisions. Trials were self-initiated by clicking on a 90 x 19 px box situated at the bottom of the screen. This step was introduced to ensure that participants would bring back the mouse to a similar starting position at the beginning of every trial. They were instructed to inspect the stimulus for as long as necessary within a limit of 6s. If they failed to answer within the 6s timeframe a buzzing sound was played and the trial was excluded from the analysis. The second part of the task consisted in giving a confidence judgement on the accuracy of the decision, on a visual analogue scale ranging from 0 (sure of having provided an incorrect response) to 100 (sure of having provided a correct response). Participants were encouraged to use the continuous scale as accurately as possible. To minimise the confounding effect of visual-discrimination task difficulty on confidence judgments, motion coherence was titrated to reach 71% of correct response for every participant using a 1-up/2-down adaptive staircase procedure preliminary to the experiment (80 trials without confidence judgments task). The staircase procedure was maintained throughout the main experiment to account for training or fatigue effects on task performance.

### sEEG data collection and preprocessing

sEEG data were collected during the course of the experiment using semi-rigid linear electrodes (Dixi Microtechniques, Besançon), with a sampling rate of 512 Hz using a Micromed recording system (Micromed, Treviso, Italy). All preprocessing steps were performed in Matlab (R2019b) using the FieldTrip toolbox (Oostenveld et al., 2011); version 20211016). Broadband signals from each channel were first visually examined to remove high levels of epileptogenic activity (17.01% of total channels). We then performed bipolar re-referencing among the remaining channels, excluding all channels that did not have a near neighbour among the artefact-free channels. Following bipolar re-referencing the broadband signal was filtered to extract high-frequency bands between 70 and 150 hz. Seven half-overlapping frequency bands were extracted from the continuous signal in steps of 20hz. Each frequency band was then split into epochs corresponding to 500 ms before stimulus onset up to the end of trials following confidence judgments. Finally, all frequency bands were baseline-corrected using the mean activity in the 500 ms preceding stimulus onset, averaged over time and trials and smoothed using a 400 ms second-order Savitzky-Golay filter. Following filtering, individual trials were visualised and trials still exhibiting important epilepsy-related artefacts were excluded (7.45% of trials were removed during these steps).

### Statistical analysis

All the analyses described in this section were performed under Python 3.8.8 in a Jupyter lab environment. All descriptive analyses were performed using Numpy and Scipy (Harris et al., 2020; Virtanen et al., 2020). All generalised linear models were fitted using StatsModel (Seabold & Perktold, 2010), and mixed generalised linear models were computed with lme4 (Bates et al., 2014) run in Python using Rpy2.

### Behavioural analyses

Trials with decision times shorter than 100 or longer than 2500 ms were excluded from the analysis (7.03% of trials), as well as trials with a change of mind (see below). To evaluate the relationship between response accuracy and decision time or confidence judgments we used mixed generalised linear models with a gamma distribution and log link function, as both decision times and confidence were positive with skewed distributions. We used the following model formula:

Decision Time ∼ Accuracy + (1|Participant) or Confidence ∼ Accuracy + (1|Participant)

### Evidence accumulation model

The model instantiates two anticorrelated accumulators, one for each possible decision outcome (Faivre et al., 2021; Kiani et al., 2014; Mazurek et al., 2003; Moreno-Bote, 2010; Pereira et al., 2020; van den Berg et al., 2016), and the confidence judgments were simulated by extending the evidence accumulation process after the initial decision. The free parameters were the rate at which sensory evidence is accumulated (drift rate), the boundary at which accumulated evidence leads to a decision (bound) and the proportion of response time that does not correspond to evidence accumulation (non-decision time). To reduce the degrees of freedom of the model, we set the covariance between accumulators to ⍴ = -0. 5, and the non-decision time standard deviation to 60 ms (van den Berg et al., 2016). Although confidence ratings were obtained on a continuous scale during the experiment, they were discretized into two bins to facilitate model fitting. We assumed that participants had high confidence when the difference between the winning and losing accumulators (Moreno-Bote, 2010; van Zandt, 2000; Vickers, 1979) *t_conf* seconds after the decision boundary exceeded a confidence threshold thr_conf.

To fit the model, we simulated 1000 trials for each participant with decision times and response accuracies. The sign of the decision time was inverted for simulated errors, allowing us to estimate the log-likelihood based on a Kolmogorov-Smirnov test between the simulated decision time distribution and those observed in the data (Pisauro et al., 2017). We fitted the free parameters with a Nealder-Mead simplex optimization procedure (Lagarias et al., 1998). To avoid local minima, we initialised the procedure with a wide range of different starting points (N = 24). To fit confidence, we used the parameters from the best decision model and simulated confidence ratings based on a readout of post-decisional evidence accumulation for 1000 trials. We computed log-likelihoods as the sum of two components. First, a binomial log-likelihood between the simulated and observed proportion of high-confidence for correct and erroneous responses. Second, a log-likelihood based on a Kolmogorov-Smirnov test between the simulated decision time distribution and those observed in the data for each confidence level and accuracy (correct or error). We optimised the confidence threshold (thr_conf) and readout time (t_conf) free parameters similarly to the decision stage, with 11 readout times ranging from 0 s post-decision (i.e. decisional readout) to 1 s post-decision. For the decisional readout, we set t_conf = 0 and only optimised thr_conf. We took the best model in terms of log-likelihood. To simulate neural data, we assumed that evidence accumulation would i) stop at the decision, ii) continue post-decisionally up to the confidence readout (Figure 5A), or for a minimum of 500 ms (Figure 5B). We also assumed that the non-decision time would be split between a pre-accumulation non-decision time of 100 ms and the remaining non-decisional time would be a post-decision non-decision time corresponding to motor delays. These assumptions did not have much influence on our conclusions.

Finally, for CoMs, we set a threshold on the difference between the winning and losing accumulators during the 2 s post-decision to reproduce the proportion of CoMs. Thus, an increase in the activity of the losing accumulator following the initial decision could bring the simulated neural activity to cross the threshold, leading to a CoM. We then derived the timing of simulated CoM from the threshold crossing times, which we compared to observed CoM timings out-of-sample to avoid overfitting.

### Trial segmentation

We focused sEEG analyses on two temporal windows of interest aimed at capturing perceptual and decisional processes. The first window was centred on stimulus onset, which was considered to precede evidence accumulation by a few hundred milliseconds. The second window was centred on mouse decision time, which we used as a proxy for decision time. Taking the decision time as a proxy for decision time rather than the time at which participants clicked on the left or right target had the advantage of removing the variability induced by different mouse trajectories between participants. All stimulus-aligned analyses were performed from 200 ms before to 1000 ms after stimulus onset and all movement-aligned analyses were performed from 500 ms before to 500 ms after decision time. Similarly to the behavioural analyses, trials with decision times inferior to 100 ms and superior to 2500 ms were excluded. Trials leading to an incorrect answer were also excluded, and trials containing changes of mind were analysed separately.

### Channel selection

Only channels in grey matter located in a subset of regions of interest were included in the analysis. These ROIs were defined according to the literature as regions likely to support evidence accumulation and/or confidence judgements (Balsdon et al., 2021; Fleming et al., 2012, 2018; Liu & Pleskac, 2011; Pereira et al., 2020, 2021). Seven ROIs were selected based on coordinates from the MarsAtlas (Auzias et al., 2016): visual cortex, parietal cortex, pre/supplementary motor cortex (pSMA), dorso-lateral prefrontal cortex (dlPFC), inferior frontal cortex (IFC), orbito-frontal cortex (OFC), and Insula (see Table 1 in SI for details). Of note, some of the deeper channels in intracranial electrodes targeting the IFC were in fact located in the insula. All channels in these electrodes were manually checked against the participants’ MRI to ensure that they were correctly labelled.

All analyses were performed on the high-gamma signal downsampled from 512 Hz to 64 Hz to decrease computation time and multiple comparisons. We additionally restricted the analysis to channels considered responsive to the stimulus. To do so, we selected channels for which the average high-gamma activity in a window covering 200 ms post-stimulus to median decision time was significantly superior to the activity during a 500 ms baseline window immediately preceding stimulus onset (trial-averaged high-gamma activity, one-sided t-test). A subset of 102 channels was found to be stimulus-sensitive across all patients.

### Single channels reflecting evidence accumulation

Channels exhibiting steeper high-gamma ramps following stimulus onset for shorter decision times were considered as reflecting evidence accumulation (Kiani et al., 2008; Kim & Shadlen, 1999). We evaluated this relationship by fitting a linear function to single trial high-gamma activity from stimulus onset to decision time, and correlating the resulting slopes with decision times (Spearman rank order correlation test). Among the 102 responsive channels, 41 were found to have a negative correlation between slopes and decision times. Permutation testing was performed by shuffling the decision times, performing the correlation again with shuffled labels and counting the resulting selected channels, giving us an estimate of the probability of finding our results by chance. This process was repeated 1000 times, and the p-value was computed as (C + 1) / (n_perms + 1) where C is the number of permutations whose score is superior or equal to the true score.

### Single channels reflecting confidence judgments

Channels reflecting confidence were selected based on the correlation between high-gamma activity in a 400 ms window preceding or following decision time and confidence judgments. The average activity in these windows of interest for each trial was correlated with confidence judgments (Pearson Rank Correlation test, threshold p-value: 0.05). Channels exhibiting a significant positive correlation between confidence and high-gamma were selected. This analysis was performed on all responsive channels, and the selected channels were labelled either mixed sensitivity channels if they had also been labelled as accumulating evidence, or pure confidence channels if it was not the case. These results were validated using permutation tests on shuffled confidence judgements, with the same procedure as described in the section above.

### Mixed-generalised linear models

We used mixed generalised linear models (mGLMs) to analyse pooled channels across participants for each ROI. Pooling the activity from several channels across participants can represent a challenge, as each participant contributes differently to the overall dataset according to idiosyncratic implantation sites. mGLMs solve this challenge by accounting for the hierarchical nature of this type of data and allowing contributions with distinct weights to the model’s predictions. We used nested mGLMs with a gamma distribution and log link function of the following forms:

Hga ∼ Decision Time + (1|Participant/channel) Hga ∼ Confidence + (1|Participant/channel).

Random intercepts were fitted for each participant and each channel within each participant. We fitted one model per data point in the 200 ms - 1000 ms window following stimulus onset and a -400 ms - 400 ms window around decision time. P-values were obtained for each time point in these windows using a Wald test procedure, which were then corrected for multiple comparisons using false discovery rate (Benjamini & Hochberg, 1995, 2000).

Additionally, we ran post hoc analyses to rule out motor confounds considering instantaneous mouse velocity Vel(t) as a covariate (see SI Table 1):

Hga ∼ Decision Time * Vel(t) + (1|Participant/channel) Hga ∼ Confidence * Vel(t) + (1|Participant/channel)

### Changes of Mind

Trials with changes of mind were analysed separately, and due to their low number, we conducted statistical analyses on all channels and trials across participants for each ROI. Participants with less than 10 changes of mind trials were excluded from the analysis (14 participants were kept). Changes of mind were detected based on crossings of the screen vertical midline occurring after at least one-third of the vertical distance to the target had been travelled. The onset of a change of mind was determined by finding the time at which the mouse cursor acceleration peaked prior to this crossing. We compared three conditions: change of mind trials aligned on the change of mind onset, regular trials aligned on decision time, and regular trials aligned on change of mind-like onset. This latter condition was to ensure that the effects observed in change of mind trials were not due to another process common to all trials. To obtain change of mind-like timing we first fitted the distribution of change of mind times for each participant with a Gaussian density function. We then randomly drew as many timings from this distribution as there were non-change of mind trials and used those timings to align the high-gamma signal. This procedure was repeated 50 times to test the robustness of our findings. To compare the effect of the different conditions over time we used a generalised linear model of the form Hga ∼ Condition with a gamma distribution and log link function. The model was applied to every time point in a -400 ms to 400 ms window, and corrected for multiple comparisons with the false-discovery rate procedure (Benjamini & Hochberg, 1995, 2000).

## Author Contributions

NF, MP developed the study concept. NF implemented experiments. DG, FS, LG, MR, NF collected data. DG, LG, MP, NF analysed data. DH performed surgeries. LM, PK, AR provided clinical data. DG, NF, MP drafted the paper. All authors provided critical revisions and approved the final version of the paper for submission.

## Acknowledgements

The authors would like to thank Clarissa Baratin, Blandine Chanteloup, and Manik Bhattacharjee for their help with the electrode localization procedure, and Emmanuelle Marmet, Marine Carmona for their help with data collection. NF has received funding from the European Research Council (ERC) under the European Union’s Horizon 2020 research and innovation programme (Grant Agreement No. 803122). Co-funded by the European Union (ERC, LEAP, 101077874). Views and opinions expressed are however those of the author(s) only and do not necessarily reflect those of the European Union or the European Research Council. Neither the European Union nor the granting authority can be held responsible for them.

## Data availability statement

Data and analysis scripts are publicly available (https://doi.org/10.17605/OSF.IO/2KT97).

The authors declare no competing interests.

## Supplementary information

### Model supplementary results

The fitted bound was 45.50 (SE = 3.59), the drift rate was 0.019 (SE = 0.002) and the non-decision time was 0.16 s (SE = 0.03). The bound for high confidence was 39.27 (SE = 3.12) and the readout time was 0.30 s (SE = 0.09) after the decision.

**Figure S1.**
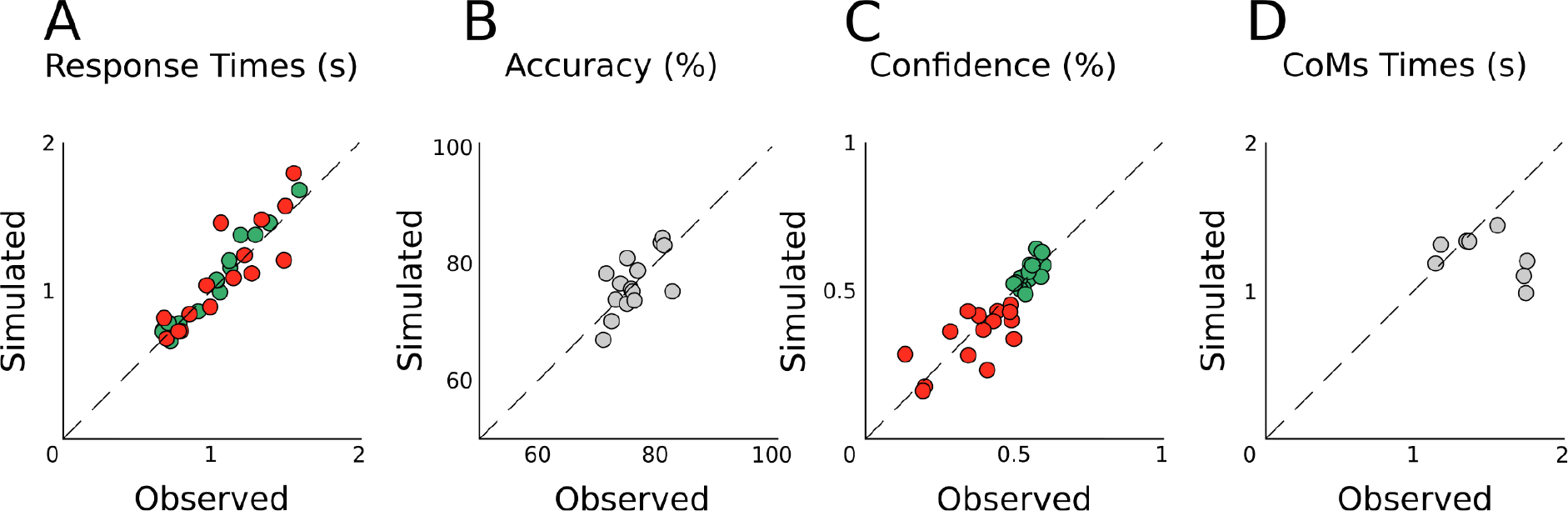
Between-participant correlation between observed and simulated data. A. Mean decision times (Spearman R = 0.96, p < 0.001 for correct trials, R = 0.90, p < 0.001 for error trials). B. mean response accuracies (R = 0.63, p = 0.014). C. Proportion of high confidence ratings (R = 0.96, p < 0.001 for correct trials, R = 0.90, p < 0.001 for error trials). D. Mean CoM timing (R = -0.24, p = 0.58)

### Model fitting with control data

We also fitted our computational model to behavioural data from 13 healthy controls (7 females, mean age = 43.5, sd = 11.7). As observed with individuals with epilepsy, the model was successful in predicting performance, decision times and confidence.

**Figure S2.**
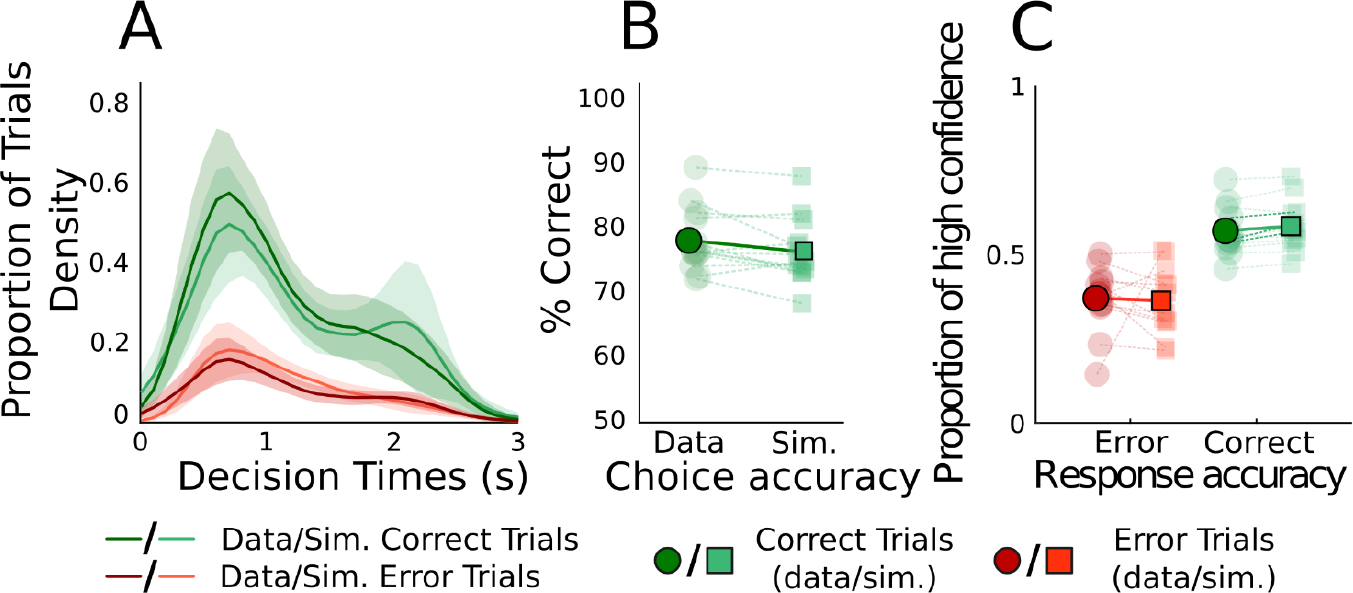
Model fits on behavioural data from healthy controls. Conventions are the same as in Figure 1. C. (**A**) Comparison of move onset timing density for data and model separated by correct and error trials (shaded areas represent 95% CI). **(B)** Choice accuracy from data and model prediction. Individual averages are plotted in light green and population averages in darker green. **(C)** Proportions of trials with low/medium/high confidence for correct and error trials comparing data and model predictions. Same convention as the middle panel.

### Regional correlates of confidence

The figure and analysis shown here are similar to Fig. 5 for the parietal and orbitofrontal cortices. We additionally display the regression coefficients, which were used for the latency analysis in Fig. 5C.

**Figure S3.**
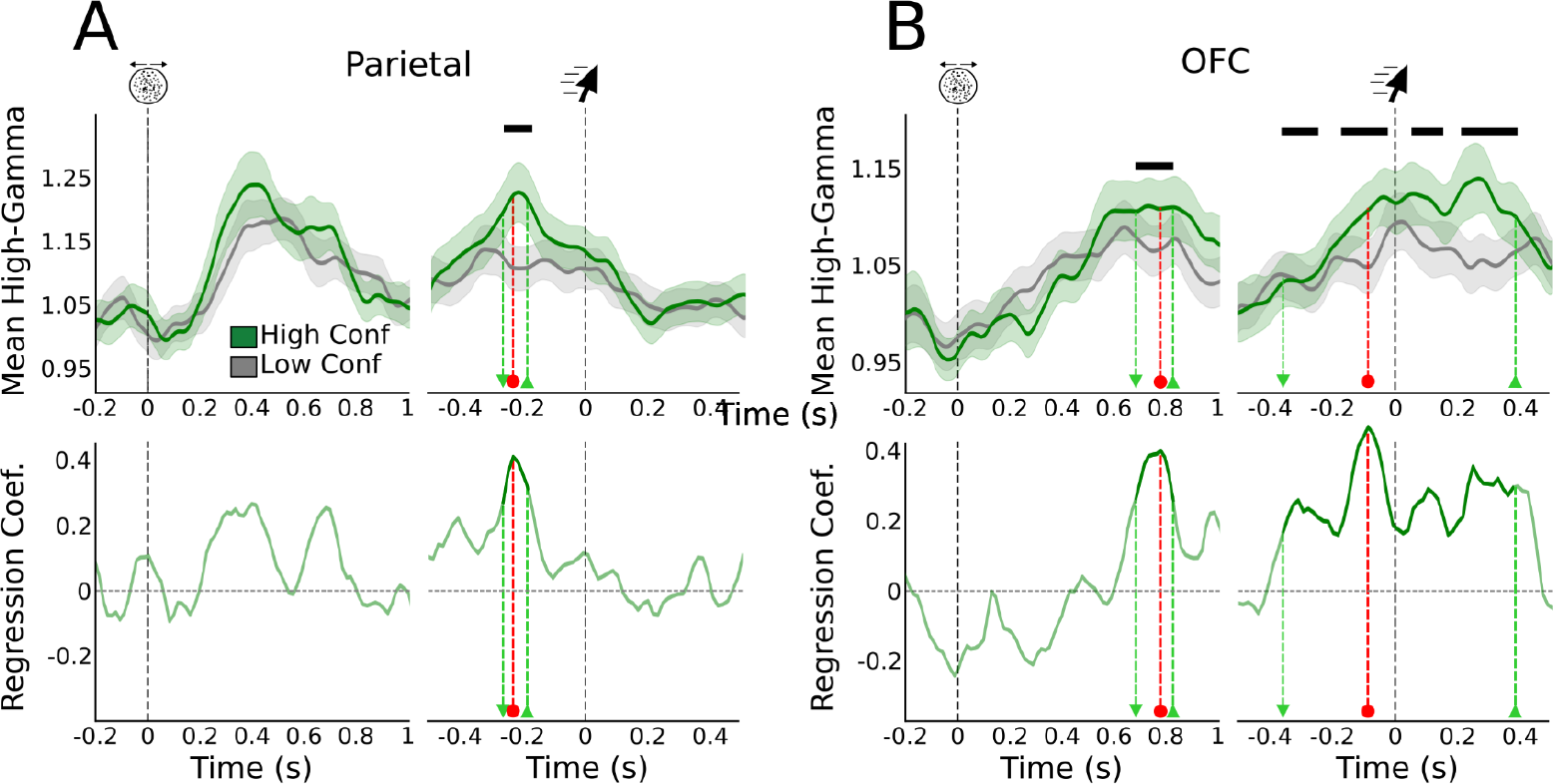
Stimulus-aligned and movement-aligned high-gamma activity for regions of interest reflecting evidence accumulation and confidence judgments (**A to B**) Top: Stimulus-aligned and movement-aligned high-gamma activity averaged across participants, including only channels reflecting evidence accumulation in each ROI for all EA channels. Dark bars indicate a significant relationship between high-gamma and confidence (p<0.05, FDR corrected using the Benjamini-Hochberg method). Green lines correspond to averaged high-gamma for the 50% of trials with the highest confidence and grey lines for the 50% lowest. Shaded areas correspond to 95% CI. Bottom: Regression coefficient as a function of time in the stimulus and movement-aligned window. Light green markers indicate the onset and offset times of significant effects, and red markers indicate peak effects.

### Performance and Confidence across nCoM and CoM trials

**Figure S4.**
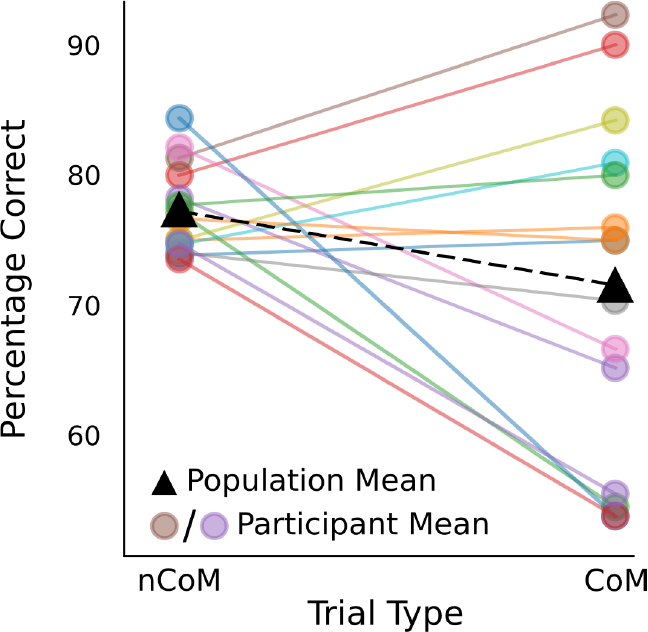
Comparison of average performance for non-change of mind and change of mind trial for each participant. Black triangles indicate population mean performance.

Changes of mind led to an overall decrease in accuracy (β = -0.306, p = 0.019), with high inter-individual variability. We also observed a significant negative effect of changes of mind on confidence (β = -4.872, p<0.01). Such diversity in the effects of CoM on accuracy and confidence has already been reported in the same task (Faivre et al., 2021).

### Regional correlates of accuracy

**Figure S5.**
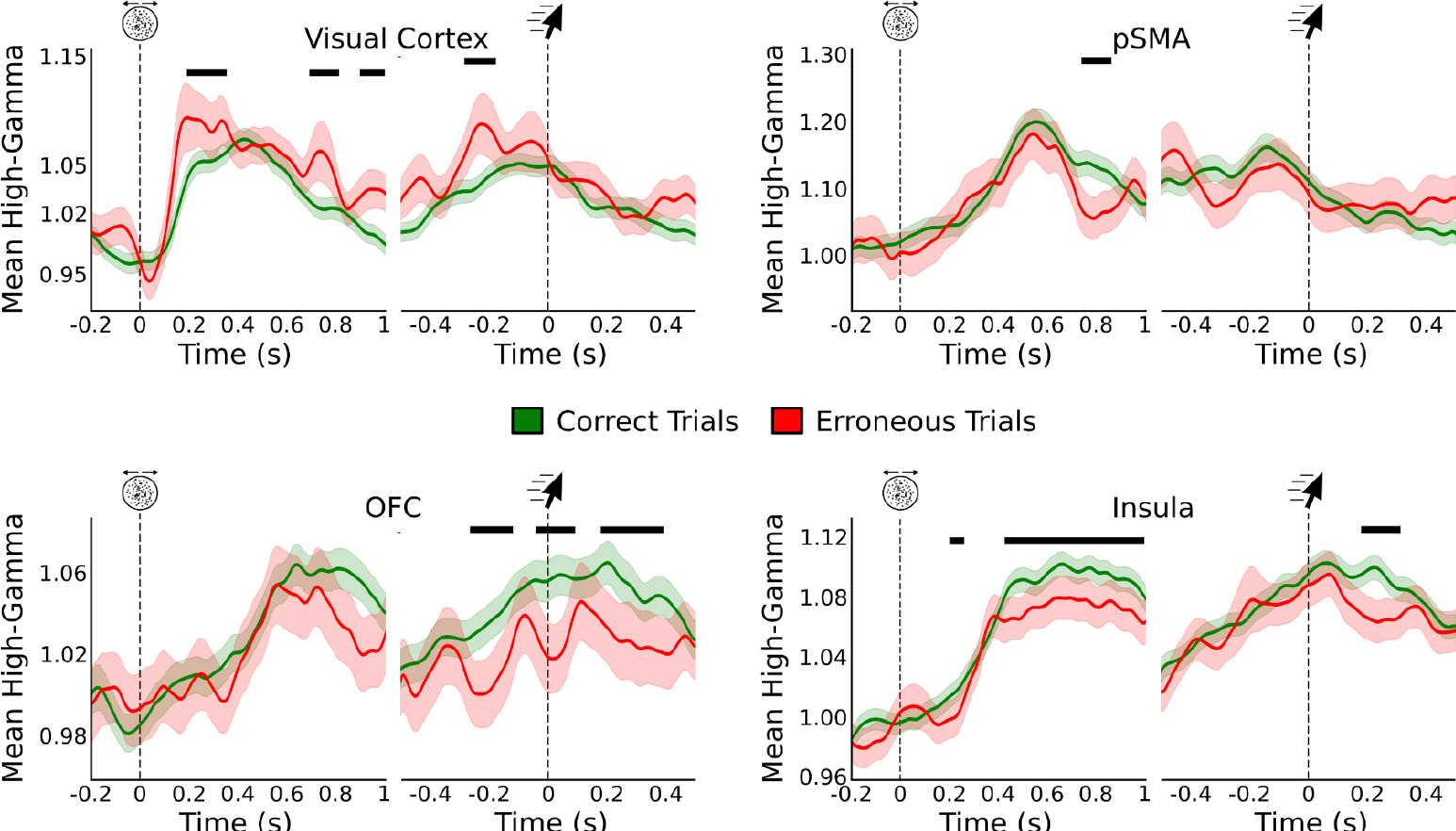
Stimulus-aligned and movement-aligned high-gamma activity averaged across participants for regions reflecting trial outcome, including all responsive channels. Dark bars indicate a significant relationship between high-gamma and trial outcome (p<0.05, FDR corrected using the Benjamini-Hochberg method). Green lines correspond to averaged high-gamma for correct trials and red lines for erroneous trials. Shaded areas correspond to 95% CI.

**Figure S6.**
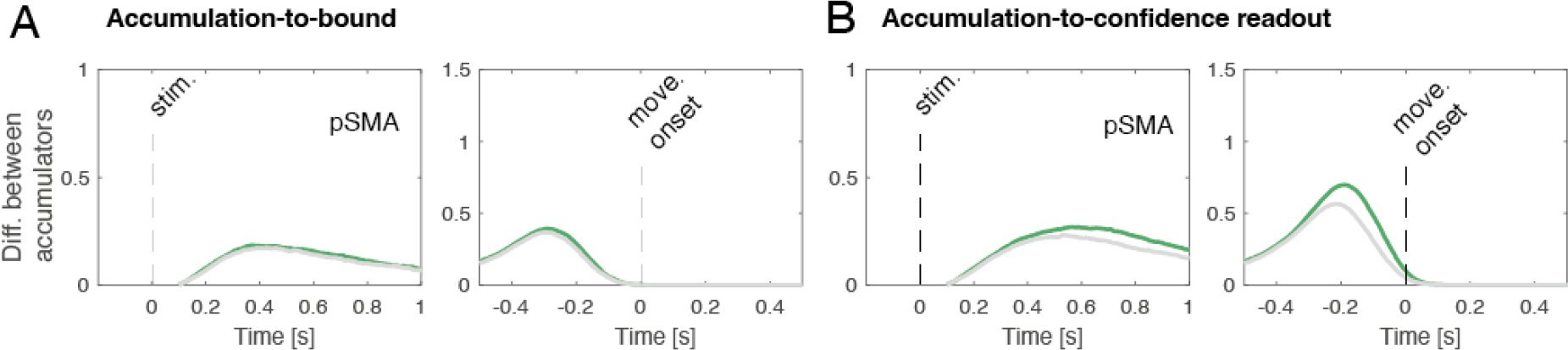
Comparison of model simulations for pSMA activity between **A.** accumulation-to-bound (accumulation stops after a decision boundary is crossed) and **B**. accumulation-to-confidence readout (accumulation continues for a few hundred milliseconds until confidence is read out) as in Figure 5A. Left panels: time window locked on stimulus onset, right panels: time window locked on movement onset.

We selected single channels sensitive to accuracy by applying a generalised model for each channel for each time point following the decision time. Only 6 of the 102 responsive channels were found to be sensitive, and none exhibited a stronger activity following incorrect decisions. Although we did not find error sensitivity with single channel analysis, we performed a region-level analysis including all responsive channels to uncover if some ROIs encoded error signals. We performed a mixed generalised linear model analysis on the activity of all responsive channels pooled within each ROI. As shown in Fig. S5, four ROIs exhibited an effect of trial outcome on accuracy. However, none of the activity in these was systematically higher for correct trials, contrary to what would have been expected for error-monitoring signals.

### Region of interest atlas

The selected ROIs were based on the Mars Atlas. The details regarding the seven ROIs as well as their corresponding Brodmann Areas (BAs) are provided below. Note that Mars Atlas-defined regions may cover only part of the associated Brodmann area.

**Table S1:** Region of interest definition

| ROI | MarsAtlas | Brodmann Areas |
| --- | --- | --- |
| **Visual cortex** | - Caudal Medial Visual Cortex (VCcm)<br>Rostral Medial Visual Cortex (VCrm),<br>Cuneus (Cu) | BA 17/18<br>BA 18/19<br>BA 18/19 |
| **Parietal cortex** | Superior Parietal Cortex (SPC)<br>Medial Superior Parietal Cortex (SPCm)<br>Dorsal Inferior Parietal Cortex (IPCd) | BA 7<br>BA 7<br>BA 39/40/7 |
| **Pre-supplementary motor area** | Dorsolateral Premotor Cortex (PMdl)<br>Dorsomedial Premotor Cortex (PMdm) | BA 6/8<br>BA 6 |
| **Dorsolateral Prefrontal Cortex** | Rostral Dorsolateral Inferior Prefrontal Cortex (Pfrdli)<br>Rostral Dorsolateral Superior Prefrontal Cortex (Pfrdls) | BA 10/46<br>BA 10/9 |
| **Inferior Frontal cortex** | Rostral Ventrolateral Prefrontal Cortex (PFrvl)<br>Rostral Ventral Premotor Cortex (PMrv) | BA 10/9<br>BA 44/45 |
| **OrbitoFrontal Cortex** | Ventral Orbito Frontal Cortex (OFCv)<br>Ventromedial Orbito Frontal Cortex (OFCvm)<br>Ventrolateral Orbito Frontal Cortex (OFCvl) | BA 11/47<br>BA 10/11<br>BA 11/47 |
| **Insula** | Insular Cortex (Insula) | BA 13 |

**Table S2:** channels repartition across ROIs and participants. Results at the ROI level are shown for simple models using decision time or confidence as a variable of interest and more complex models where instantaneous mouse velocity is added as a covariate.

| ROI | Resp. channels | EA channels | Conf. channels | Mixed channels | ROI level EA | ROI level EA (Vel. Model) | ROI level Conf. | ROI level Conf. (Vel. Model) | ROI level CoM |
| --- | --- | --- | --- | --- | --- | --- | --- | --- | --- |
| Visual Cortex | 17 (N = 3) | 8 (N = 2) | 2 (N = 1) | 2 (N = 1) | No | No | No | No | No |
| Parietal Cortex | 8 (N = 3) | 4 (N = 3) | 3 (N = 1) | 1 (N = 1) | No | No | Yes | Yes | No |
| pSMA | 9 (N = 4) | 7 (N = 3) | 2 (N = 1) | 2 (N = 1) | Yes | Yes | Yes | Yes | Yes |
| dIPFC | 10 (N = 6) | 1 (N = 1) | 1 (N = 1) | 0 (N = 0) | No | No | No | No | Yes |
| IFC | 7 (N = 3) | 6 (N = 4) | 0 (N = 0) | 0 (N = 0) | Yes | Yes | No | No | No |
| OFC | 17 (N = 7) | 4 (N = 4) | 6 (N = 4) | 3 (N = 3) | No | No | Yes | Yes | Yes |
| Insula | 34 (N = 12) | 11 (N = 5) | 14 (N = 6) | 7 (N = 3) | Yes | Yes | Yes | Yes | Yes |
| **Total** | **102** | **41** | **28** | **15** |  |  |  |  |  |

### Analysis of channels outside ROIs

To ensure that our ROIs selection did not skew our results we performed an exploratory analysis including all channels that were not included in the main article. We performed the same steps as described in the main text, namely selecting channels that were responsive to the stimulus, and among them identifying channels reflecting evidence accumulation and/or confidence. Of the total of 724 channels initially excluded from our analysis 69 were found to be responsive to the stimulus (9.53% of included channels, against 24.63% for the main text analysis). Among these, 31 channels were found to accumulate evidence, a number that significantly exceeded what was expected by chance (permutation test, p = 0.0009). Similarly, among the 69 responsive channels, 3 were found to reflect confidence, failing permutation testing (permutation test, p >0.01). Considering that no significant cluster of channels exhibiting signatures of evidence accumulation or confidence emerged, we did not perform analyses at the level of ROIs.

### Relationship between neural markers of confidence and decision time

Considering the correlation between decision times and confidence, we considered the possibility that the positive relationship we observed between high-gamma activity and confidence judgments may be partly driven by decision times. Although it is not possible to fully disambiguate the effects of decision times and confidence on the channels reflecting confidence, we could measure the respective strength of these effects and compare them. We did so by computing the correlation between high-gamma activity in a 400 ms window following the move onset and decision times and confidence. We reasoned that, if our effects were fully driven by decision times, the magnitude of the correlation with decision times should be stronger than with confidence. We found on the contrary that in this window the high-gamma levels correlated more strongly with confidence than decision times, confirming that the effect we observe is at the very least partly driven by confidence levels. The same results were found when considering the channels reflecting confidence in a 400ms pre-decisional window.

We also tested this potential confound at the population level by running a mixed-effects regression for confidence while including an interaction term with the following formula:

Hga ∼ Confidence + Confidence:Decision time + (1|sub/channel).

The effect of confidence was preserved in all ROI, indicating that the interaction between confidence and decision time does not account for the effect of confidence on high-gamma activity.

**Table S3:** Fraction of channels removed during preprocessing

| sub | Total channels | Removed | Remaining | Proportion remaining (%) |
| --- | --- | --- | --- | --- |
| 1 | 122 | 3 | 119 | 97.54 |
| 2 | 122 | 0 | 122 | 100 |
| 3 | 122 | 14 | 108 | 88.52 |
| 4 | 122 | 53 | 69 | 56.55 |
| 5 | 122 | 19 | 103 | 84.42 |
| 6 | 122 | 20 | 102 | 83.60 |
| 7 | 122 | 59 | 63 | 51.63 |
| 8 | 122 | 4 | 118 | 96.72 |
| 9 | 122 | 0 | 122 | 100 |
| 10 | 122 | 6 | 116 | 95.08 |
| 12 | 122 | 15 | 107 | 87.70 |
| 13 | 122 | 11 | 111 | 90.9 |
| 14 | 122 | 8 | 114 | 93.44 |
| 16 | 184 | 29 | 155 | 84.23 |
| 17 | 143 | 28 | 115 | 80.41 |
| 18 | 136 | 58 | 78 | 57.35 |
| 19 | 169 | 7 | 162 | 95.85 |
| 20 | 204 | 104 | 100 | 49.01 |
| 21 | 186 | 32 | 154 | 82.79 |
| 22 | 185 | 25 | 160 | 86.48 |
| 23 | 122 | 1 | 121 | 99.18 |
| **Total** | **2915** | **496** | **2419** |  |
| **Mean** | **138.80** | **23.61** | **115.19** | **83.88** |
| **Std** | **27.87** | **25.98** | **26.80** | **16.24** |

**Table S4:** Participants’ demographics and antiseizure medication. Antiseizure medication corresponds to the daily dose prescription.

| Participant | Age | Sex | Handedness | Antiseizure medication |
| --- | --- | --- | --- | --- |
| 1 | 41 | M | R | Clobazam 15mg<br>Lacosamide 300mg<br>Oxcarbazepine 1200mg |
| 2 | 25 | F | L | Clobazam 10mg<br>Lacosamide 400mg<br>Perampanel 6mg |
| 3 | 28 | F | R | Carbamazepine 400mg<br>Felbamate 3000mg |
| 4 | 44 | F | R | Carbamazepine 400mg<br>Lamotrigine 200mg<br>Levetiracetam 1500mg |
| 5 | 46 | M | R | Brivaracetam 200mg<br>Clobazam 10mg<br>Lamotrigine 300mg<br>Perampanel 8mg |
| 6 | 18 | M | R | Carbamazepine 1200mg<br>Zonisamide 200mg |
| 7 | 25 | F | R | Brivaracetam 400mg<br>Lacosamide 350mg<br>Zonisamide 200mg |
| 8 | 34 | F | R | Clobazam 5mg<br>Lacosamide 400mg<br>Levetiracetam 1750mg<br>Perampanel 10mg |
| 9 | 20 | M | R | Carbamazepine 1300mg<br>Clobazam 5mg<br>Lacosamide 200mg |
| 10 | 52 | M | R | Carbamazepine 800mg<br>Lacosamide 400mg |
| 12 | 28 | M | R | Carbamazepine 1200mg<br>Levetiracetam 2250mg |
| 13 | 37 | F | L | Carbamazepine 800mg<br>Clobazam 5mg<br>Levetiracetam 2000mg |
| 14 | 21 | M | R | Carbamazepine 1200mg<br>Clobazam 20mg<br>Perampanel 6mg |
| 16 | 28 | M | R | Clobazam 10mg<br>Eslicarbazepine acetate 1200mg<br>Zonisamide 400mg |
| 17 | 39 | M | R | Lacosamide 600mg<br>Perampanel 6mg |
| 18 | 24 | F | L | Lacosamide 300mg<br>Lamotrigine 300mg |
| 19 | 37 | M | R | Carbamazepine 1200mg |
| 20 | 21 | M | R | Carbamazepine 1200mg<br>Clobazam 40mg<br>Zonisamide 150mg |
| **21** | 45 | F | L | Carbamazepine 1200mg<br>Clonazepam 1mg<br>Levetiracetam 3000mg<br>Topiramate 300mg |
| **22** | 27 | M | R | Carbamazepine 800mg<br>Clobazam 5mg<br>Levetiracetam 2000mg |
| **23** | 30 | M | R | Carbamazepine 800mg<br>Clobazam 15mg<br>Perampanel 10mg |

